# The Metabolic Organ Clock: A Computable Framework Linking Cell-Lineage Architecture to Emergent Organ Bioenergetics

**DOI:** 10.64898/2026.09.29.755490

**Authors:** Alejandra Martin Sevilla, Lurong Pan

## Abstract

Cell-type atlases now catalogue hundreds of distinct human cell identities across development, each annotated with mitochondrial abundance, dominant fuel pathway, and lineage of origin, yet no computable model connects the topology of the lineage tree to the metabolic rate the resulting organs display in the adult body. Using a curated 396-node human cell-lineage tree spanning the zygote to terminal somatic identities, we test whether organ-level standard metabolic rate (SMR) is better predicted by developmental time (lineage depth) or by terminal fate identity (anatomical compartment). We find that lineage depth explains essentially none of the variance in a cell’s metabolic tier (r = 0.11, R^2^ ≈ 1.2%), whereas compartment identity explains roughly three-quarters of it (≈75%), and that the mean mitochondrial volume fraction of an organ’s constituent terminal cell types tracks the organ’s classical literature-derived SMR with r = 0.90 across five canonical reference-man organ groups. These results motivate a five-layer computable framework — lineage as a formal tree grammar, information-theoretic structure of the tree, metabolic switches as runtime operators, compilation to organ-level energetic networks, and execution as organism-level bioenergetics — and a specific falsifiable hypothesis we call the metabolic commitment-horizon model: that a cell’s metabolic tier is fixed at a discrete lineage-commitment event and thereafter held constant, rather than accumulated continuously with differentiation time. In a worked held-out test (5-fold cross-validation, n = 396), a nearest-commitment-ancestor predictor beat a global-mean null (MAE 0.752 vs 0.809, p = 0.015) but was not significantly better than a lineage-depth regression (ΔMAE = −0.042, 95% CI −0.098 to +0.012, p = 0.175), so we report the commitment-horizon model as supported descriptively but not yet discriminated predictively on these data. We then connect this within-species result to the classical, cross-species rate-of-living hypothesis, in which breathing rate scales predictably with body mass (Stahl’s allometric equation, fR = 53.5·M^−0·26^) yet fails to predict lifespan on its own: naked mole rats, little brown bats, humans, and bowhead whales all live several-fold longer than mass alone predicts (longevity quotients of 4.4, 5.1, 4.6, and 2.8 respectively), for documented reasons unrelated to a slower pace of living. We then treat insulin resistance as an acquired perturbation of the clock, and find that insulin responsiveness is orthogonal to mitochondrial density: adipose and skeletal muscle, the two lowest-mitochondrial-density tissues in the organ set, are the principal sites of insulin-stimulated glucose disposal, while brain, the second-highest by per-kilogram rate, is essentially insulin-independent. We survey machine learning and deep learning tools — graph neural networks, single-cell foundation models (Geneformer, scGPT, scVI), and RNA velocity — already available to extend or directly test this framework on primary single-cell data. We report the supporting computations transparently, flag which values in the underlying dataset are measured versus estimated or drawn from the literature, and close with eight testable predictions and their falsification conditions.

## 1. Introduction

Single-cell atlases and developmental biology together have produced detailed maps of which cell types exist, when they arise, and what they are made of, but comparatively little computable machinery connects a cell’s position in the lineage tree to the energetic behavior of the organ it ultimately builds. Epigenetic “clocks” such as Horvath’s multi-tissue DNA-methylation predictor recover chronological age from methylation state with remarkable accuracy [1], demonstrating that a cell’s molecular profile can encode a computable notion of biological time. No comparable clock exists for cellular bioenergetics: it is not established whether a cell’s metabolic phenotype — its mitochondrial volume fraction, its dominant fuel pathway, its position on a low-to-high metabolic-tier scale — is a function of how many differentiation steps separate it from the zygote, or whether it is instead fixed by which terminal fate it is committed to, independent of path length.

This distinction matters because it determines what kind of model is appropriate. If metabolic phenotype accumulates gradually with developmental time, a continuous, depth-indexed model (analogous to a molecular clock) is the right object to build. If instead metabolic phenotype is set at discrete commitment events and held constant thereafter, the right object is a piecewise, event-indexed model — closer to a state machine than a clock in the colloquial sense. We use the word “clock” throughout in this second, computational sense: a rule that outputs a cell’s or organ’s metabolic state as a function of its position in a lineage structure, not a predictor of chronological or biological age.

### 1.1 Summary of new results

Because the manuscript is organized as a layered framework, we state the original computational results up front so they are not obscured by the scaffolding. All five are derived here from the dataset described in the Methods, and all five are new.

i. **Fate identity dominates developmental time**. Anatomical compartment explains approximately 75% of the variance in cellular metabolic tier, whereas lineage depth explains 1.2% (Section 3.2). The depth relationship is statistically significant (p = 0.030) yet negligible in magnitude, a dissociation we report explicitly.
ii. **Cellular mitochondrial content recovers organ-level metabolic rate**. Mean mitochondrial volume fraction of an organ’s constituent cell types correlates with its classical reference SMR at r = 0.90 across five organ groups (Section 5.1).
iii. **Metabolic switching is decoupled from lineage branching**. Across 131 internal nodes, a node’s branching factor is uncorrelated with the fraction of its outgoing edges that change tier (r = −0.015), indicating that fate specification and metabolic switching are separable programs (Section 4.2).
iv. **A held-out test of the commitment-horizon model returns an equivocal result**. Under 5-fold cross-validation, a nearest-commitment-ancestor predictor beats a global-mean null (p = 0.015) but is not significantly better than a lineage-depth regression (p = 0.175). We report this negative finding rather than the descriptive statistic that appears to favour the model (Section 6.4).
v. **Insulin responsiveness is orthogonal to mitochondrial density**. The two lowest-mitochondrial-density tissues in the organ set, adipose and skeletal muscle, are the principal sites of insulin-stimulated glucose disposal, while brain is essentially insulin-independent — an axis the mitochondrial-tier model does not capture (Section 8.2).

### 1.2 A layered computational frame, applied to lineage instead of sequence

We adopt the general strategy of decomposing a biological object into layers, each with an established formalism, and asking where the interfaces between layers are the most productive and most fragile. Here the base object is not a genome sequence but a rooted, 396-node human cell-lineage tree running from the zygote to terminal somatic identities, annotated with germ layer, anatomical compartment, developmental depth, mitochondrial volume percentage, dominant fuel pathway, a five-level ordinal metabolic tier (T0–T5, non-respiring to very high), and, for a subset of terminal cell types, the literature-derived standard metabolic rate (SMR, kcal·kg^−1^·day^−1^) of the organ they compose. The five layers we propose are: (1) lineage as a formal tree grammar, (2) information-theoretic structure of the tree, (3) metabolic switches as runtime operators along lineage edges, (4) compilation from cellular composition to organ-level energetic networks, and (5) execution as organism-level bioenergetics — the layer at which the commitment-horizon clock hypothesis lives. Sections 2–6 develop each layer in turn with results computed directly from the dataset; Section 7 connects the within-species result to the cross-species rate-of-living literature on breathing rate and longevity; Section 8 treats insulin resistance as an acquired perturbation of the clock and shows that hormonal responsiveness is an axis the mitochondrial-tier model does not capture; Section 9 surveys machine learning and deep learning tools within reach for extending or testing the framework, layer by layer; Section 10 states the framework’s limits without euphemism; Section 11 closes with eight falsifiable predictions.

**Figure 1.**
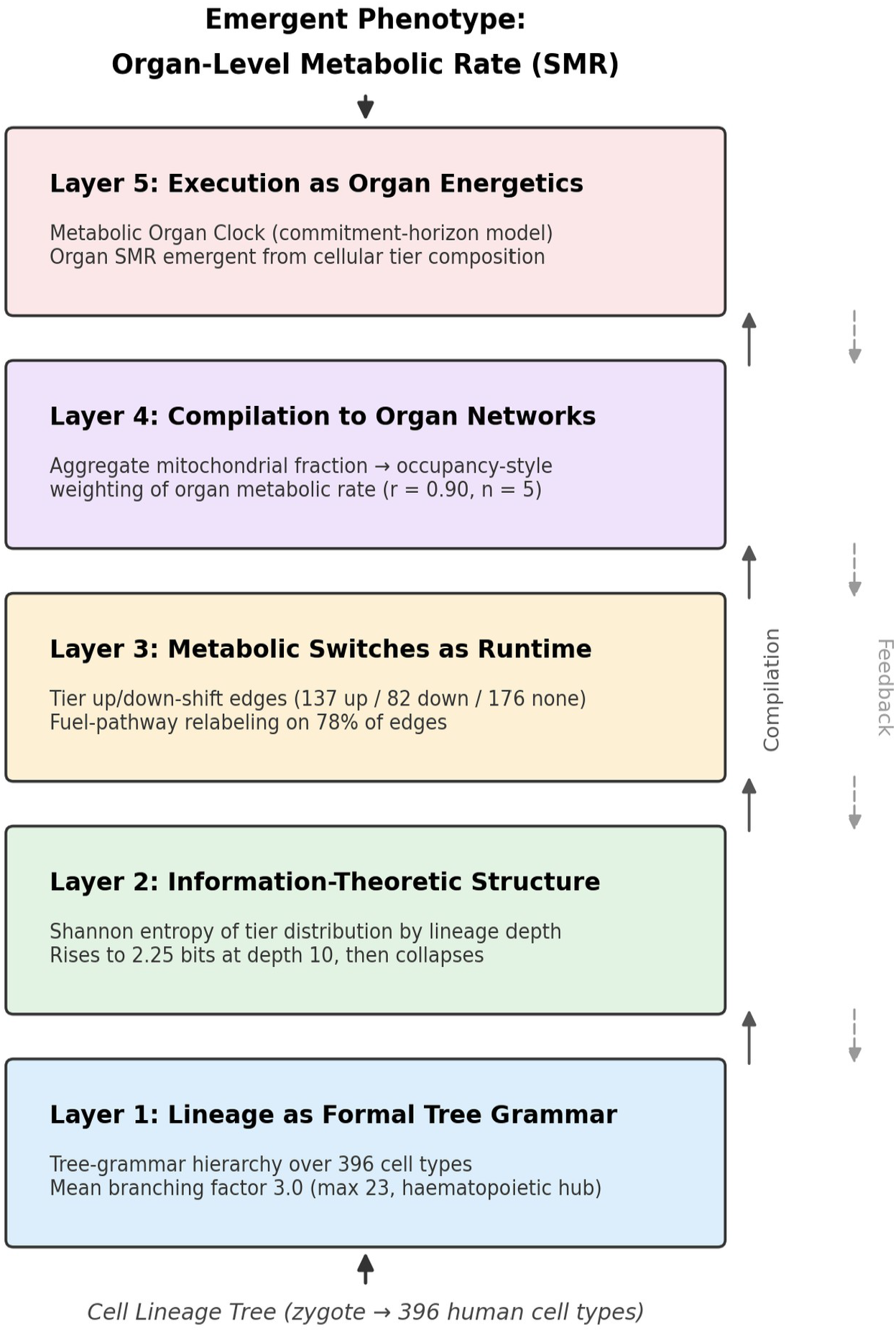
The five-layer computable framework proposed in this paper, instantiated with the quantitative results reported in Sections 2–6. Solid arrows denote compilation from lower to higher layers; dashed arrows denote feedback/validation paths a full framework would need to close.

## 2. Layer 1: Lineage as a Formal Tree Grammar

A lineage tree is naturally described by a tree grammar: production rules that rewrite a parent cell-type symbol into an ordered or unordered set of child symbols. Tree-automata theory [2] provides the same kind of hierarchy for tree languages that the Chomsky hierarchy provides for string languages, and mapping the observed lineage tree onto it clarifies which parts of development are describable by simple, local rules and which require history-dependent or context-dependent rewriting.

### 2.1 Branching structure of the tree

Across the 131 internal (non-terminal) nodes in the dataset, the mean branching factor is 3.0 children per parent (median 2, maximum 23). A regular tree grammar — one production per symbol, no dependence on ancestral history beyond the immediate parent — is sufficient to describe the majority of bifurcating or trifurcating transitions in the tree (for example, the zygote-to-blastomere-to-trophectoderm/inner-cell-mass split). It is not sufficient at the small number of high-branching hub nodes: the single largest branch point in the dataset is a haematopoietic-type progenitor with 23 direct descendants, which requires either a context-free tree grammar with reusable non-terminal categories (myeloid, lymphoid, erythroid sub-programs invoked repeatedly) or an explicitly context-sensitive rule in which the same progenitor symbol rewrites differently depending on niche signals not encoded in the tree topology itself — exactly the situation described for chromatin-level regulation in sequence-based computable-genome frameworks, where the acting alphabet is not the linear structure but an external contact or signaling graph.

### 2.2 A tree-grammar hierarchy for lineage

**Table 1.** A tree-grammar hierarchy applied to the 396-node lineage dataset, by analogy with the Chomsky hierarchy applied to genomic sequence elements.

| Grammar class | Lineage feature | Example from the dataset |
| --- | --- | --- |
| Regular tree grammar | Local, history-independent bifurcation | Zygote → Blastomere → Trophectoderm / Inner cell mass |
| Context-free tree grammar | Reusable sub-programs invoked at multiple points | Repeated deployment of a generic “epithelial → basal/suprabasal” motif across skin, gut, and airway lineages |
| Context-sensitive rewriting (claimed) | Fate depends on niche signal external to tree topology | Neural crest sub-populations (cranial, cardiac, trunk, vagal) diverging into germ-layer-crossing fates by target tissue |
| High-branching hub nodes | Single parent, many direct fates (n up to 23) | Haematopoietic-type progenitor giving rise to myeloid, lymphoid, and erythroid descendants |

As with the sequence case, the honest statement is narrower than the framing device suggests: very few of the context-sensitive-looking transitions in this dataset are formally context-sensitive tree rewrites. Most are regular or context-free rewrites with an unencoded niche variable standing in the background. The tree-grammar layer is useful for locating where that hidden variable must live, not for eliminating it.

## 3. Layer 2: Information-Theoretic Structure of the Lineage Tree

Independent of any specific grammar, the lineage tree can be treated as a labeled structure whose information content is measured directly. We compute the Shannon entropy of the five-level metabolic tier (T0–T5) among all cell types sharing a given lineage depth, and separately decompose the total variance in cellular metabolic tier into a component explained by anatomical compartment (terminal fate identity) and a residual component.

### 3.1 Tier entropy rises with depth, then collapses

**Figure 2.**
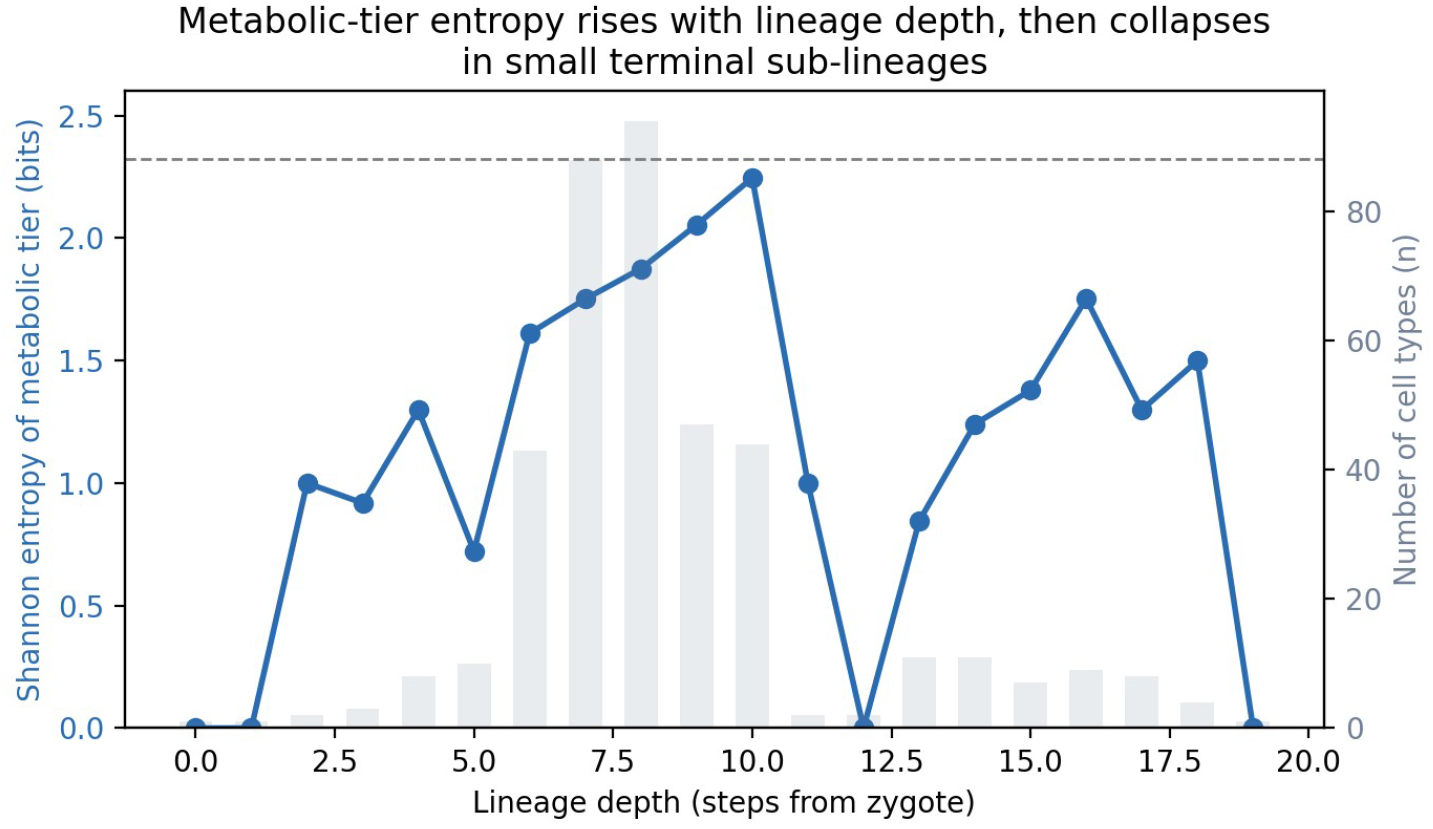
Shannon entropy (bits) of the metabolic-tier distribution among cell types sharing a given lineage depth (blue line, left axis), against the number of cell types sampled at that depth (grey bars, right axis). The dashed line marks the maximum possible entropy for five tiers (log_2_ 5 ≈ 2.32 bits).

Entropy rises from 0 bits at depths 0–1 (the zygote and its immediate descendants are metabolically uniform by construction) to a peak of 2.25 bits at depth 10, close to the theoretical ceiling of 2.32 bits for a five-way categorical variable. This is the signature expected under progressive fate restriction: as the tree branches, the population of cell types at a given depth becomes an increasingly even mixture of low- and high-tier fates. Entropy then collapses toward depths 11–19, but this second regime should not be over-interpreted: cell-type counts drop sharply beyond depth 10 (from dozens per depth to single digits), and the deep tail of the tree is dominated by a small number of narrow, highly specialized clades (skin appendage and eye lineages reach depth 15–18), so the apparent collapse is at least partly a small-sample artifact rather than a biological reconvergence of metabolic state.

### 3.2 Fate identity, not depth, is where the information lives

A stronger and more direct test asks how much of the total variance in cellular metabolic tier is explained by anatomical compartment (a proxy for terminal fate identity) versus by lineage depth (a proxy for developmental time). Across 227 distinct compartments, compartment identity explains approximately 75% of the total variance in metabolic tier (pooled within-compartment variance / total variance ≈ 0.25, so between-compartment variance ≈ 0.75). Lineage depth alone, by contrast, explains only about 1.2% of the variance in tier. The same near-null relationship holds between depth and mitochondrial volume fraction directly (r = −0.055), while tier and mitochondrial volume fraction are, as expected from their shared construction, strongly correlated with each other. The information-theoretic layer therefore points in a single direction: metabolic state is compressible almost entirely by knowing what a cell is going to become, not by knowing how many divisions or specification steps it took to get there.

**Table 1b.** Ordinary least squares regression statistics for the two bivariate relationships underlying Section 3.2’s variance decomposition.

| Relationship | n | Slope (SE) | R<sup>2</sup> | p-value |
| --- | --- | --- | --- | --- |
| Tier ~ Depth | 396 | 0.035 (0.016) | 0.012 | 0.030 |
| Mitochondrial % ~ Tier | 395 | 4.70 (0.20) | 0.576 | < 0.001 |

The tier-versus-depth regression is a useful illustration of why effect size and statistical significance are not the same question: with n = 396, a slope as small as 0.035 tiers per lineage step is nonetheless distinguishable from zero (p = 0.030), yet it accounts for barely one-hundredth of the variance in tier (R^2^ = 0.012). A relationship can be statistically significant and practically negligible at the same time, and the depth-versus-fate contrast in this paper rests on the effect size, not on significance testing. The mitochondrial-percentage-versus-tier regression, by contrast, is both significant and substantial (R^2^ = 0.576), consistent with tier having been assigned from mitochondrial content in the first place rather than being an independent measurement.

**Figure 2b.**
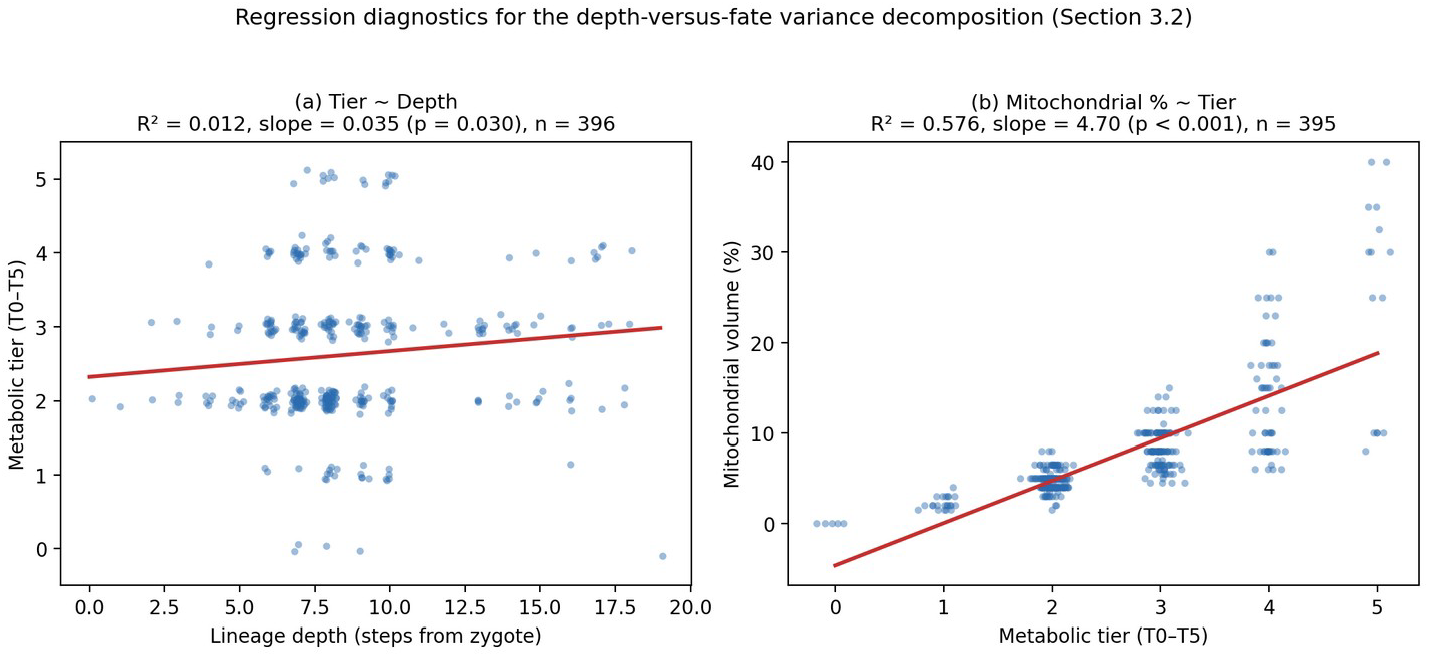
Regression diagnostics for both relationships in Table 1b. (a) Metabolic tier against lineage depth: points are visibly scattered across the full tier range at every depth, and the fitted line (red) is nearly flat — the regression is significant only because of the large sample size, not because depth has real predictive value. (b) Mitochondrial volume percentage against metabolic tier: the fitted line tracks a clear, substantial increasing trend, consistent with tier having been assigned from mitochondrial content directly. Points jittered slightly in (a) and (b) to reduce overplotting on the discrete tier axis.

## 4. Layer 3: Metabolic Switches as Runtime Operators

If fate, not depth, carries the metabolic information, the interesting computational objects are the specific lineage edges at which metabolic state changes — the “instructions” that move a cell from one tier or fuel program to another. We treat every parent→child edge in the tree as a candidate switch event and classify it by the direction of tier change.

### 4.1 Switch statistics

Of 395 parent→child lineage edges, 137 (35%) show a tier increase, 82 (21%) show a tier decrease, and 176 (45%) show no tier change. A description of the dominant fuel/pathway field changes across 308 edges (78%), reflecting both genuine metabolic reprogramming and finer-grained relabeling within an unchanged tier. Two well-characterized biological programs recovered by this classification give the down-shift edges external validation: lens epithelial cells losing most of their organelles, including mitochondria, as they terminally differentiate into lens fibre cells to achieve optical transparency [3], and keratinocytes progressively down-regulating oxidative demand through the basal→spinous→granular→corneocyte sequence as they cornify. Both are real, independently documented programmed-organelle-clearance events, not artifacts of the tier encoding, and both appear in the dataset as consecutive tier-decreasing edges.

### 4.2 Branching hubs are not preferentially switch-rich

A natural hypothesis is that high-branching hub nodes — the ones requiring context-free or context-sensitive grammar in Section 2 — are also where the largest fraction of metabolic switching happens, since these are the points where the widest range of fates is being specified. Across the 131 internal nodes, the correlation between a parent’s branching factor and the fraction of its outgoing edges that change tier is effectively zero (r = −0.015). Branching multiplicity and metabolic-switch frequency are, in this dataset, independent properties of a lineage node: some very high-branching hubs (the haematopoietic-type progenitor) preserve tier across most descendants, while several low-branching or unary transitions (e.g., single-child terminal differentiation steps) carry the largest tier changes. This is a negative result worth stating plainly rather than working around.

**Table 2.** Lineage-edge transition statistics treated as runtime switch operators, analogous to mobile-element or state-transition operators in sequence-based computable frameworks.

| Switch type | Count / fraction | Representative example |
| --- | --- | --- |
| Tier increase | 137 / 395 (35%) | Progenitor → metabolically demanding terminal cell (e.g., cardiomyocyte lineage) |
| Tier decrease | 82 / 395 (21%) | Lens epithelial cell → lens fibre cell (organelle clearance); keratinocyte cornification |
| No tier change | 176 / 395 (45%) | Sequential differentiation steps within one metabolic band |
| Fuel-pathway relabeling | 308 / 395 (78%) | Glycolysis-leaning → mixed → OXPHOS-dominant along a single branch |

## 5. Layer 4: Compilation — From Cellular Composition to Organ-Level Energetics

The fourth layer asks whether a static property of a lineage’s terminal cell types — here, mitochondrial volume fraction — compiles into the emergent, whole-organ energetic behavior that physiology has measured independently. This is the layer at which the framework is most directly testable, because organ-level SMR has a long, largely settled measurement literature to check against.

### 5.1 An occupancy-style aggregation model

Elia’s widely used reference-man organ metabolic-rate table [4], subsequently evaluated and confirmed by whole-body indirect-calorimetry modeling in adults [5], assigns standard metabolic rates of 440 kcal·kg^−1^·day^−1^ to heart and kidney, 240 to brain, 200 to liver, 13 to skeletal muscle, and 4.5 to adipose tissue. The dataset used here assigns these same five literature values to the relevant terminal cell types by anatomical compartment. Aggregating the dataset’s independently curated mitochondrial-volume-percentage field by these five organ groups gives mean mitochondrial volume fractions of 13.9% (heart/kidney), 7.6% (brain), 9.6% (liver), 4.75% (skeletal muscle), and 3.0% (adipose).

**Table 2b.**
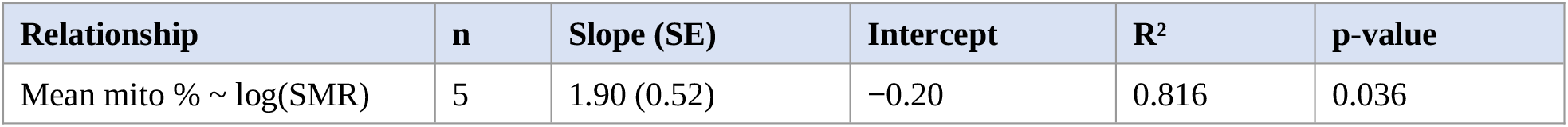
Ordinary least squares regression of mean mitochondrial volume fraction on log(organ SMR), across the five organ anchor groups shown in Figure 3.

The fitted line (mean mitochondrial % ≈ −0.20 + 1.90·ln(SMR)) reaches conventional significance despite the minimal sample size (p = 0.036, n = 5, 3 degrees of freedom), and explains 82% of the variance in mitochondrial fraction across the five organ groups. Given the small n, this p-value should be read as descriptive rather than confirmatory — it quantifies how tight this particular five-point fit is, not the probability that the underlying biological relationship is real across organs in general, which is a separate question addressed by Prediction 2 in Section 10.

**Figure 3.**
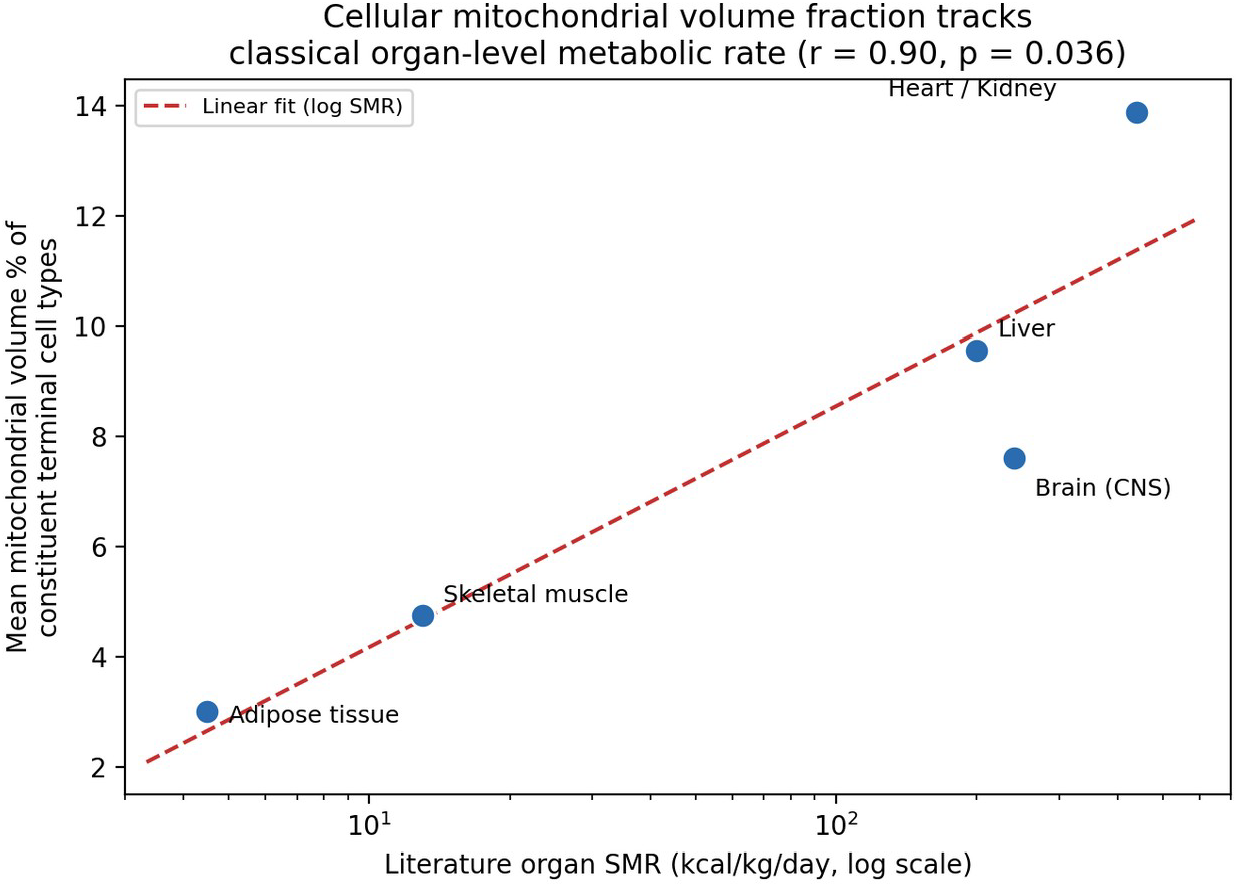
Mean mitochondrial volume percentage of an organ’s constituent terminal cell types, plotted against the organ’s classical reference-man standard metabolic rate [4,5] (log scale). n = 5 organ groups.

This is the layer-4 analogue of a thermodynamic occupancy model: rather than deriving gene expression from a Boltzmann-weighted sum over promoter binding configurations, we are deriving organ energetic output from an (unweighted, cell-number-agnostic) average over the mitochondrial density of the cell types an organ is built from. The fact that a cellular ultrastructural feature curated independently of any organ-level measurement recovers the classical between-organ ordering of metabolic rate is, at minimum, a non-trivial consistency check on both the dataset and the general principle — well established in cellular bioenergetics [6] — that oxidative capacity scales with mitochondrial content at the tissue level.

### 5.2 A necessary caveat on independence

The five SMR anchor values are not five independent measurements: they are drawn from a single classical source (Elia, 1992 [4], subsequently re-evaluated by Wang et al., 2010 [5]), and the mitochondrial-volume-percentage field, while curated separately, was almost certainly written with the same qualitative expectation (heart and kidney very high, adipose very low) in mind. The r = 0.90 correlation should be read as “the dataset’s independently entered cellular annotations reproduce a well-known physiological ordering,” which is a genuine and useful validation of the layer-4 aggregation logic, but it is not a five-fold-replicated discovery, and it does not yet demonstrate that the same aggregation would predict the SMR of an organ not already anchored to one of these five categories. Section 11 turns this into an explicit, falsifiable out-of-sample prediction.

## 6. Layer 5: Execution — The Metabolic Commitment-Horizon Clock

The final layer asks what kind of “clock” the results of Sections 3–5 actually license. Two candidate models are available. A continuous clock would predict a cell’s metabolic tier as a smooth, monotonic function of lineage depth — the metabolic analogue of a methylation-based age predictor, where the signal accumulates steadily with developmental time [1]. A commitment-horizon clock would instead predict tier as a step function of fate: fixed at the lineage node where a cell commits to a terminal or near-terminal identity, and held constant along every subsequent edge within that identity, regardless of how many further morphological or transcriptional steps remain.

### 6.1 The depth model fails; the fate model does not

The evidence from Sections 3 and 4 favors the second model specifically. Depth explains 1.2% of tier variance; compartment identity explains roughly 75%. Branching multiplicity, which would be expected to correlate with switch frequency under a continuous-accumulation model (more branch points passed, more opportunities for gradual metabolic drift), shows no such relationship (r = −0.015). And the switch-edge analysis in Section 4 shows metabolic tier changing in sharp, large steps at specific, biologically identifiable commitment points (organelle clearance in lens fibre and keratinocyte lineages) rather than drifting gradually across many small edges.

### 6.2 A formal statement of the model

We define the commitment-horizon model formally as follows. For a lineage tree with tier-labeling function T, let a node v be a commitment node if the subtree rooted at v is tier-homogeneous (all descendants of v share v’s tier) while v’s parent’s subtree is not. The model predicts that for any node u, T(u) = T(c(u)), where c(u) is the nearest commitment-node ancestor of u — that is, tier is inherited from the most recent commitment event, not accumulated as a function of depth(u) − depth(c(u)). This is a piecewise-constant, event-indexed clock rather than a continuous, time-indexed one, and it is directly falsifiable: it predicts zero residual correlation between tier and within-clade depth after conditioning on the identity of the nearest commitment ancestor, a prediction we did not exhaustively test at the single-clade level in this analysis and flag as the immediate next computational step.

### 6.3 Relation to epigenetic age clocks

This model is deliberately not an aging clock. Epigenetic clocks such as Horvath’s [1] predict chronological or biological age from a tissue’s current methylation state and are explicitly time-indexed — methylation drift accumulates continuously after a cell is born. The metabolic commitment-horizon clock proposed here instead predicts a cell’s steady-state metabolic tier from its fate identity, and is explicitly non-time-indexed within a committed lineage. The two are complementary rather than competing: an aging clock could in principle be layered on top of a commitment-horizon baseline to capture drift away from a fate’s nominal tier with chronological age or metabolic disease, but that combined model is outside the scope of the present, purely developmental dataset.

### 6.4 A worked test: held-out prediction of metabolic tier

The preceding subsections argue for the commitment-horizon model from variance decomposition and edge statistics, both of which are descriptive summaries of the whole dataset. A stronger test is predictive: if the model is right, knowing a cell’s nearest labelled ancestor should predict its tier better than knowing how many steps it sits from the zygote. Because this is directly computable from the supplied lineage tree, we ran it rather than proposing it.

We compared four predictors of metabolic tier under 5-fold cross-validation over all 396 tier-labelled nodes.

(i) A global-mean null model. (ii) A depth regression: ordinary least squares on lineage depth, fitted on the training fold. (iii) A compartment-mean model, assigning each held-out node the mean tier of its anatomical compartment in the training fold, standing for fate identity. (iv) A nearest-ancestor model operationalizing the commitment-horizon hypothesis of Section 6.2: each held-out node is assigned the tier of its closest ancestor that appears in the training fold, walking the lineage tree rootward. Critically, the commitment structure is never read off the test nodes themselves; ancestors are only consulted if they were observed during training, so the comparison is genuinely out-of-sample. Errors are reported as mean absolute error (MAE) in tier units, with 95% confidence intervals from 2,000 bootstrap resamples and paired Wilcoxon signed-rank tests on per-node absolute errors.

**Table 3.** Cross-validated held-out prediction of metabolic tier (5-fold, n = 396). Lower MAE is better. ΔMAE is the paired difference against the depth regression; negative values favour the listed model.

| Model | MAE | 95% CI | ΔMAE vs depth | p (paired) |
| --- | --- | --- | --- | --- |
| Global mean (null) | 0.809 | 0.757–0.864 | +0.016 | 0.0003 |
| Depth regression | 0.793 | 0.740–0.851 | — | — |
| Nearest ancestor (commitment-horizon) | 0.752 | 0.674–0.828 | −0.042 | 0.175 |
| Compartment mean (fate identity) | 0.728 | 0.667–0.789 | −0.065 | 0.010 |

**Figure 4.**
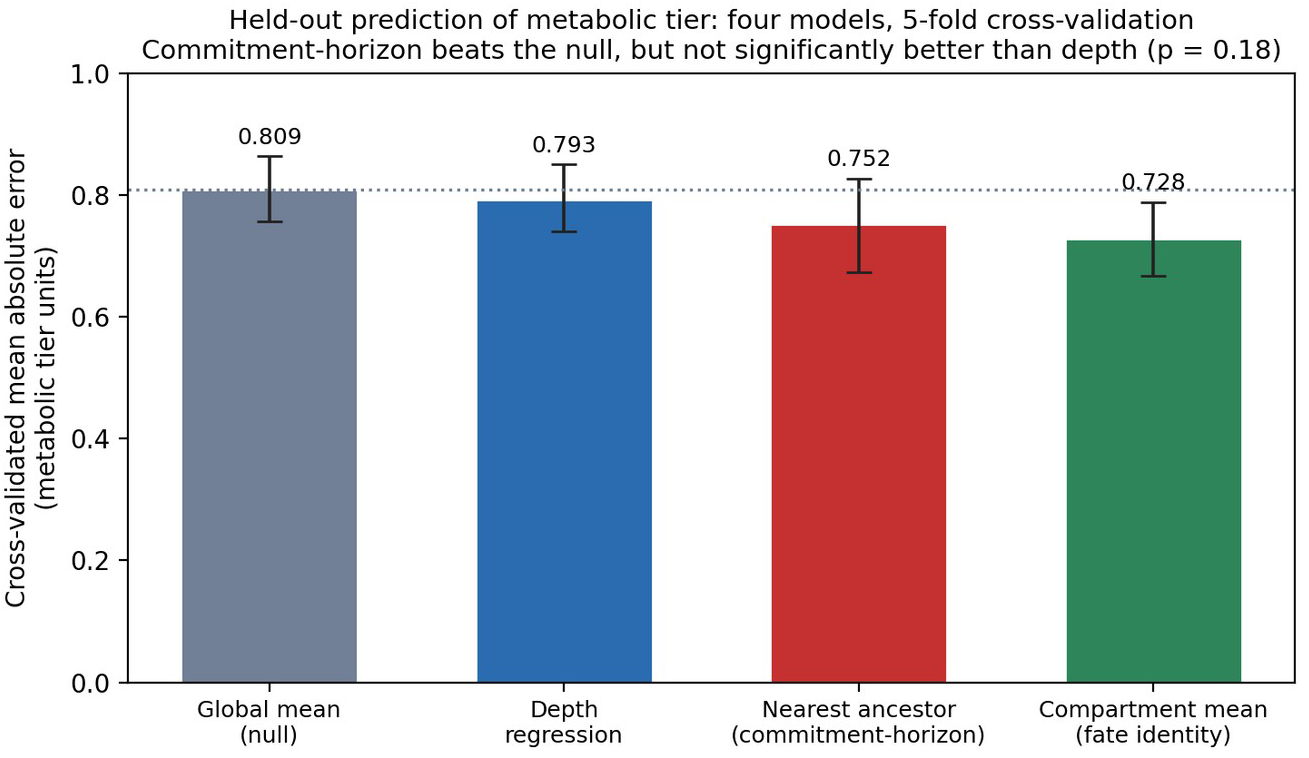
Cross-validated mean absolute error for four predictors of metabolic tier. Error bars are 95% bootstrap confidence intervals; the dotted line marks the null-model MAE. The commitment-horizon predictor improves on the null, but its advantage over the depth regression is within confidence bounds.

The result is partially, but not fully, consistent with the commitment-horizon model, and we report it as such. The nearest-ancestor predictor does beat the null model (MAE 0.752 vs 0.809; Δ = −0.057, p = 0.015), confirming that lineage position carries real information about metabolic tier. It also has a lower point-estimate error than the depth regression, in the direction the model predicts. But that specific advantage — the comparison Prediction 4 actually rests on — is not statistically significant (ΔMAE = −0.042, 95% CI −0.098 to +0.012, p = 0.175). On this dataset, we cannot claim that the commitment-horizon predictor out-predicts a continuous depth model; we can only say it is not worse and trends better.

Two further observations sharpen the picture. First, the compartment-mean model is the best of the four (MAE 0.728) and is indistinguishable from the nearest-ancestor model (Δ = −0.024, p = 0.747). Fate identity and nearest-commitment-ancestor are, predictively, the same information here — which is what the commitment-horizon model would expect if commitment nodes largely coincide with compartment boundaries, but which also means the tree structure adds nothing beyond the compartment label. Second, the depth regression does significantly beat the null (Δ = −0.016, p = 0.0003) despite explaining only 1.2% of variance (Section 3.2), reinforcing the point made there that significance and effect size come apart at this sample size.

The honest summary is that the descriptive variance decomposition of Section 3.2 overstates how cleanly this dataset separates the two hypotheses. A ∼75%/1% variance split sounds decisive; under held-out prediction the same data yield a 0.06-tier MAE gap between the best fate-based model and the null, and no significant separation between the commitment-horizon and depth predictors. Prediction 4 in Section 11 is accordingly restated as partially tested rather than confirmed, and the discriminating experiment it calls for — on primary single-cell data with finer-grained metabolic readouts than a five-level ordinal tier — remains necessary.

## 7. From One Species to Many: Rate-of-Living Theory and Its Exceptions

Sections 2–6 stayed within a single species and asked whether a cell’s metabolic state is a function of developmental time or of fate. A much older literature asks the analogous question one level up: across species, is an organism’s pace of life — its breathing rate, heart rate, metabolic rate — a predictor of how long it lives? This is the “rate-of-living” hypothesis, and breathing rate is its most literal instantiation: an animal that breathes faster is, on this view, spending its fixed lifetime allotment of oxidative damage more quickly. It is worth walking through explicitly, because it fails in the same qualitative way the continuous-depth clock failed in Section 6 — the naive time-indexed story is not the one the data support — and because the reason it fails is instructive for how the commitment-horizon idea generalizes.

### 7.1 The classical hypothesis and a real scaling law

Rubner first proposed, in 1908, that mammals expend a roughly similar amount of metabolic energy per gram of tissue over a lifetime, so that species living fast (high mass-specific metabolic rate) simply die young, and species living slow die old [8]. Breathing rate is one of the cleanest measured correlates of this pace: Stahl’s classical allometric analysis of mammalian respiratory variables gives resting breathing frequency as fR = 53.5·M^−0^·^26^ breaths/min, with body mass M in kilograms [9]. This single equation, fit originally on the rat-to-human size range, predicts a roughly 60-fold drop in breathing rate from an 8-gram bat to a 60-tonne bowhead whale. It is important to be explicit about what Figure 5 does and does not show. For most of the ten species we have not compiled an independent measurement, and plotting the equation’s own output at those masses would demonstrate nothing: a curve cannot validate itself. Those species are therefore drawn as open symbols marking equation values, not data. For the four species where standard reference ranges are available — mouse, dog, human, and horse [29] — the measured resting rates are plotted with their ranges. These scatter around the allometry rather than lying on it, falling between 0.79× (human, 14 vs 17.7 predicted) and 1.32× (horse, 14 vs 10.6 predicted) of the predicted value, with mouse at 1.17× (163 vs 140) and dog at 0.81× (20 vs 24.6). Agreement within roughly ±30% across four orders of magnitude of body mass is respectable for an allometric law, but it is a weaker claim than a figure of points lying exactly on a line would suggest.

**Figure 5.**
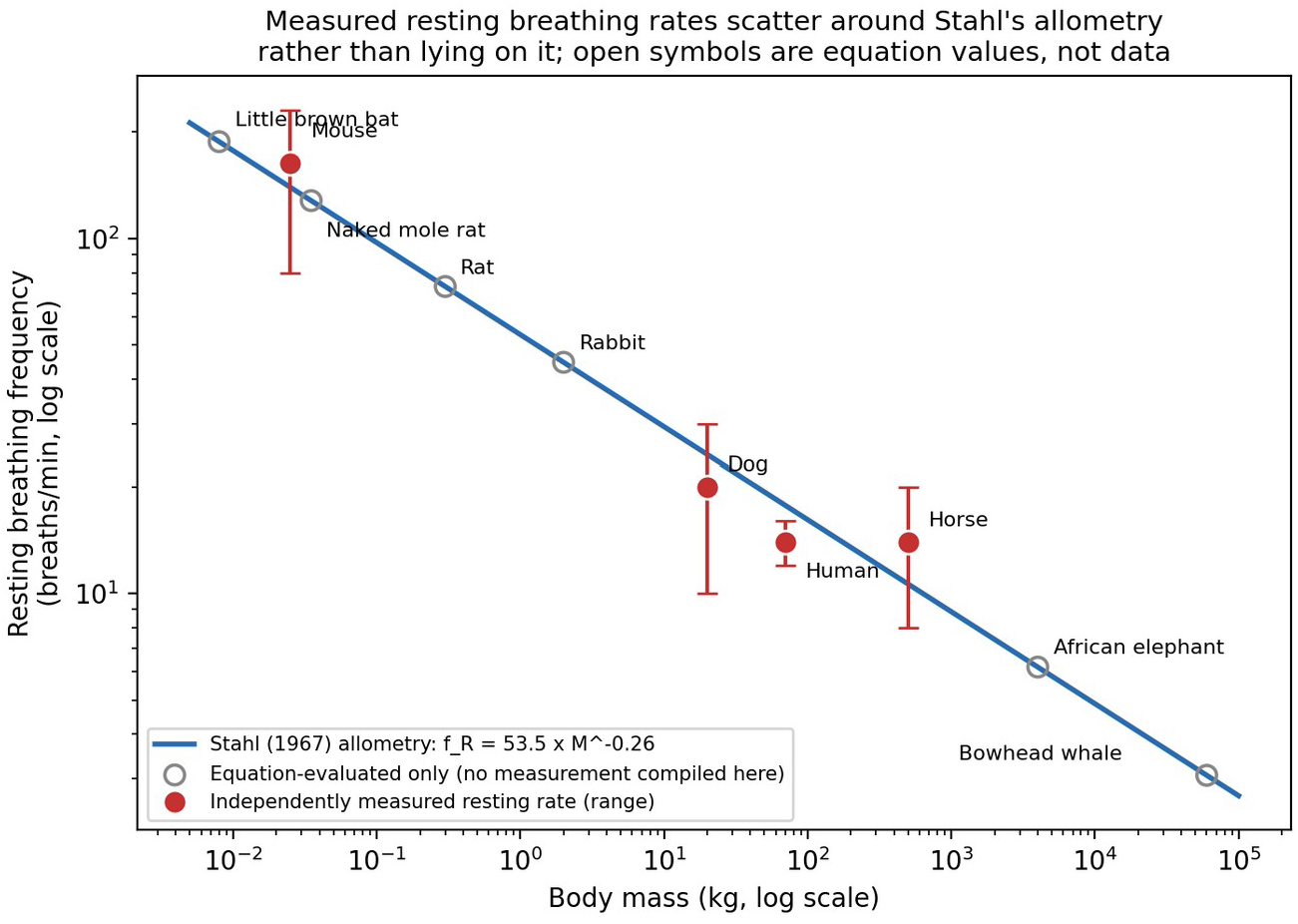
Resting breathing frequency against body mass for ten mammals. The line is Stahl’s allometric equation [9]. Filled red points with bars are independently measured resting rates and their reference ranges [29]; open grey circles are the equation evaluated at that species’ mass, included to locate the species on the axis and explicitly not evidence for the relationship. Measured values scatter between 0.79× and 1.32× the prediction. The bowhead whale point is shown for scale only: cetaceans are intermittent breathers whose surface respiration is not the same measurement as continuous resting ventilation in a terrestrial mammal.

### 7.2 The strong form of the theory fails, predictably

If breathing rate tracked lifetime oxidative exposure in a simple, fixed way, maximum lifespan should be almost fully predictable from body mass alone, since mass sets breathing rate so tightly. Using de Magalhães, Costa, and Church’s allometric lifespan equation — maximum lifespan (years) ≈ 4.88·(mass in grams)^0^·^153^, fit across the AnAge comparative-longevity database [10] — as the mass-only prediction, four of the ten species in Figure 6 live several times longer than their body mass predicts: the little brown bat (34 observed vs. 6.7 predicted years, longevity quotient LQ = 5.1), the naked mole rat (37 vs. 8.4 years, LQ = 4.4), humans (122.5 vs. 26.9 years, LQ = 4.6), and the bowhead whale (an estimated 211 vs. 75.6 years, LQ = 2.8). This is not a fringe reading of the data: de Magalhães and colleagues’ own phylogenetically corrected analysis found that basal metabolic rate does not correlate with longevity in eutherian mammals once body mass and phylogeny are properly accounted for [10], directly contradicting the strong form of the rate-of-living hypothesis on its own comparative terms.

**Figure 6.**
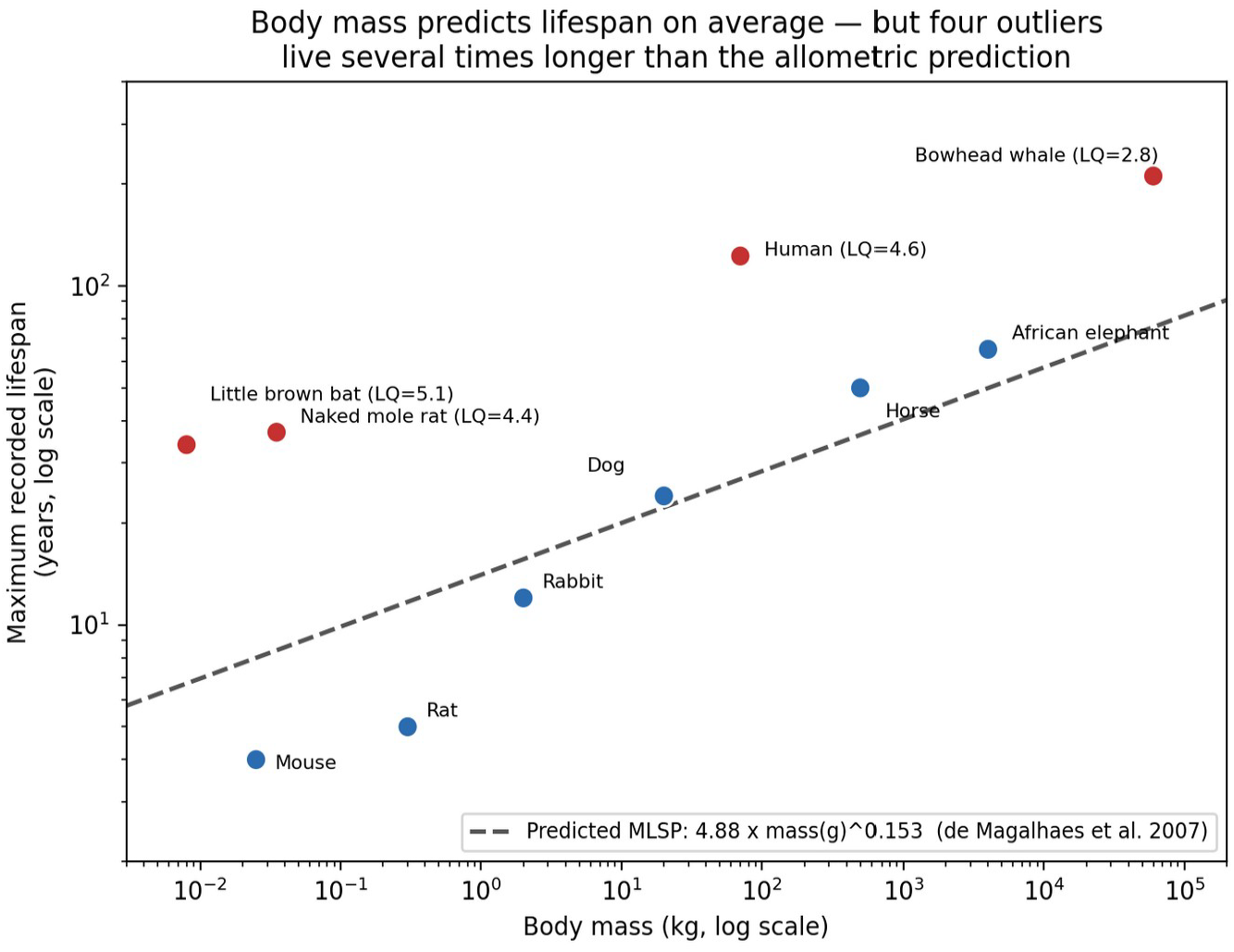
Maximum recorded lifespan against body mass for the same ten mammals, with the allometric prediction of de Magalhães et al. [10]. Longevity quotients (LQ = observed / predicted) are shown for the four species living more than 2.5× longer than mass alone predicts.

**Table 4.** Breathing rate (Stahl 1967 [9]), observed maximum lifespan (MLSP), mass-only predicted MLSP (de Magalhães et al. 2007 [10]), and longevity quotient for ten mammals. Body masses and observed MLSPs are representative literature values [10–14]; the naked mole rat LQ of 4.4 matches the published value exactly [11], validating the calculation against the source literature.

| Species | Mass (kg) | fR (breaths/min) | Observed MLSP (yr) | Predicted MLSP (yr) | LQ |
| --- | --- | --- | --- | --- | --- |
| Little brown bat | 0.008 | 187.7 | 34.0 | 6.7 | 5.07 |
| Naked mole rat | 0.035 | 127.9 | 37.0 | 8.4 | 4.40 |
| Mouse | 0.025 | 139.6 | 4.0 | 8.0 | 0.50 |
| Rat | 0.30 | 73.2 | 5.0 | 11.7 | 0.43 |
| Rabbit | 2.0 | 44.7 | 12.0 | 15.6 | 0.77 |
| Dog | 20.0 | 24.6 | 24.0 | 22.2 | 1.08 |
| Human | 70.0 | 17.7 | 122.5 | 26.9 | 4.55 |
| Horse | 500.0 | 10.6 | 50.0 | 36.3 | 1.38 |
| African elephant | 4000.0 | 6.2 | 65.0 | 49.9 | 1.30 |
| Bowhead whale | 60000.0 | 3.1 | 211.0 | 75.6 | 2.79 |

### 7.3 Why the outliers break the pattern

In each of the four outlier cases, the literature attributes the extra lifespan not to a lower pace of living — bats and naked mole rats do not breathe unusually slowly for their mass; Figure 5 places them exactly where Stahl’s equation predicts — but to a mechanism that decouples oxidative throughput from oxidative damage. Little brown bat mitochondria produce one-half to one-third the hydrogen peroxide per unit of oxygen consumed compared with two similarly sized, short-lived mammals (a shrew and a white-footed mouse), despite indistinguishable superoxide dismutase activity, directly supporting a damage-mitigation rather than a pace-reduction explanation [11]. Naked mole rats combine a metabolic rate already somewhat below allometric expectation with an essentially flat age-specific mortality hazard — in a large captive demographic study, the risk of death did not increase with age even 25-fold past reproductive maturity, a pattern inconsistent with simple cumulative oxidative wear [13]. Bowhead whales combine very large body size (which alone predicts long life) with cold body temperature and a hypothesized enhanced DNA damage response, though the aspartic-acid-racemization age estimates that place their maximum lifespan near or above 200 years carry substantial uncertainty and should be read as an order-of-magnitude claim rather than a precise figure [14]. Human longevity is the one outlier in this set that is not primarily physiological: it reflects culture, medicine, and social buffering layered on top of a primate metabolic rate, not a distinct cellular damage-suppression mechanism, and should not be read as evidence for the same class of adaptation as the other three.

### 7.4 Connecting back to the commitment-horizon clock

Sections 6 and 7 are different analyses at different scales — one within a single human lineage tree, one across ten mammalian species — and this paper does not claim they are the same phenomenon. But they share a structure worth naming explicitly. Within the human lineage tree, the naive story (“metabolic state accumulates with developmental time”) fails, and what actually predicts a cell’s metabolic tier is which fixed fate it occupies. Across mammalian species, the naive story (“lifespan is set by pace of living, and pace of living is set by size”) also fails, and what predicts the four major exceptions is a species-specific, evolved property (reduced radical production, flat mortality hazard, cold body temperature) layered on top of, not derived from, body-size-determined breathing rate. In both cases the raw rate variable — lineage depth in one, breathing frequency in the other — is a poor predictor once the right categorical variable (fate identity; species-specific damage-suppression biology) is accounted for. Section 11 turns the parallel into an explicit, falsifiable prediction rather than leaving it as an analogy.

### 7.5 Scope: why this section is mammals only, and what that choice hides

Stahl’s equation is a mammalian allometry, and every species in Table 4 is an endotherm. That restriction is not innocuous, and it is worth stating plainly what it excludes, because the most extreme longevity records in the vertebrates belong to animals the analysis above cannot accommodate. A Galapagos tortoise (Chelonoidis niger) reaches roughly 150–175 years in captivity, and the Greenland shark (Somniosus microcephalus) is the longest-lived vertebrate known, with radiocarbon dating of eye-lens nuclei placing one individual at 392 ± 120 years [30]. Neither can be placed on Figure 5: an ectotherm has no mass-determined resting breathing frequency in the sense Stahl’s equation describes, because its metabolic rate is set jointly by body mass and ambient temperature rather than by mass alone, and a tortoise’s ventilation is intermittent and posture-dependent in ways a mammalian resting rate is not.

The substantive point is that this exclusion is not neutral with respect to the hypothesis being tested. In both of these cases the longevity is conventionally attributed precisely to a low mass-specific metabolic rate, itself a consequence of low and stable body temperature — cold Arctic water for the Greenland shark, ectothermy for the tortoise. That is the rate-of-living hypothesis apparently succeeding, in exactly the clades that Section 7.2’s mammal-only analysis leaves out. The four mammalian outliers of Figure 6, by contrast, achieve longevity without a reduced pace of living, through damage suppression (Section 7.3). Read together, the two observations suggest that what actually generalizes is not “slow metabolism extends life” but something narrower: body temperature and oxidative damage rate are the operative variables, and metabolic rate correlates with lifespan only insofar as it tracks them. Restricting the comparison to endotherms, all of which hold body temperature nearly constant, removes the single variable that most plausibly drives the cross-clade pattern. A version of this analysis extended across ectotherms and endotherms together, with body temperature as an explicit covariate, would be a considerably stronger test than the one presented here, and we do not claim to have performed it.

## 8. Perturbing the Clock: Insulin Resistance as an Acquired Metabolic State

Everything to this point has treated a cell’s metabolic state as something set during development and then held. Section 6 argued specifically that tier is fixed at a commitment event and inherited unchanged thereafter; Section 7 argued that across species, longevity outliers achieve their advantage through damage suppression rather than through a slower resting pace. Neither argument addresses the obvious clinical fact that adult metabolic phenotype is not actually static. Insulin resistance — the progressive failure of skeletal muscle, adipose tissue, and liver to respond normally to circulating insulin — develops within a single lifetime, in cells whose developmental lineage and fate identity have long since been fixed. It is the cleanest available test case for what the commitment-horizon model does and does not claim, and working it through exposes a real gap in the framework as presented.

### 8.1 What insulin resistance does not contradict

The commitment-horizon model as stated in Section 6.2 is a claim about steady-state tier assignment along a lineage tree, not a claim that a committed cell’s metabolic behavior is immune to regulation. A myocyte in an insulin-resistant individual is still a myocyte: its mitochondrial volume fraction, its fate identity, and its position in the lineage tree are unchanged, and its nominal metabolic tier in the sense used throughout this paper is unchanged. What changes is its responsiveness to a hormonal signal — specifically, the insulin-stimulated translocation of GLUT4 (SLC2A4) transporter vesicles to the plasma membrane, the final common effector of insulin-stimulated glucose uptake in muscle, adipose, and cardiac tissue [28]. Insulin resistance is therefore best understood in this framework as a Layer 4 phenomenon: a change in how a compiled regulatory network responds to an input, not a change in the Layer 1–3 lineage assignment that produced the cell. The commitment-horizon clock survives, but only because it was never making a claim about hormonal regulation in the first place — which is precisely the limitation this section is written to expose.

### 8.2 Insulin responsiveness is orthogonal to metabolic tier

The more interesting result is quantitative. Insulin’s glucoregulatory action is strongly tissue-partitioned: skeletal muscle alone accounts for roughly 70–80% of insulin-stimulated glucose disposal under hyperinsulinemic-euglycemic clamp conditions [26,27], the method introduced by DeFronzo, Tobin, and Andres to quantify insulin sensitivity directly [25]. GLUT4 is the predominant insulin-responsive transporter in skeletal muscle, cardiac muscle, and adipose tissue; liver instead responds to insulin chiefly through suppression of hepatic glucose production rather than GLUT4-mediated uptake; and brain glucose uptake proceeds largely through the insulin-independent GLUT1 and GLUT3 transporters [27,28]. Mapping this partition onto the same five organ anchors used in Section 5 produces a dissociation that the mitochondrial-tier clock does not predict.

**Table 5.** Insulin-responsiveness overlay on the organ anchors of Section 5. Per-kg SMR and mean mitochondrial volume percentage are as in Table 2b and Figure 3; organ masses are standard reference-adult values and absolute energy use is their product. Transporter and insulin-class assignments are from the insulin-physiology literature [25–28], not from the cell-lineage dataset.

| Organ | Transporter | Insulin class | SMR (kcal/kg/d) | Mito % | kcal/day |
| --- | --- | --- | --- | --- | --- |
| Skeletal muscle | GLUT4 | Responsive | 13 | 4.75 | 364 |
| Adipose tissue | GLUT4 | Responsive | 4.5 | 3.00 | 68 |
| Heart | GLUT4 | Responsive | 440 | 13.88 | 145 |
| Liver | GLUT2 | Modulated | 200 | 9.56 | 360 |
| Brain (CNS) | GLUT1/3 | Independent | 240 | 7.61 | 336 |
| Kidneys | GLUT1/2 | Independent | 440 | 13.88 | 136 |

The two tissues with the lowest mitochondrial volume fraction in the entire anchor set — adipose (3.0%) and skeletal muscle (4.75%) — are precisely the two principal sites of insulin-stimulated glucose disposal, and together they account for about three-quarters of the absolute resting energy flux residing in GLUT4-expressing tissue. Conversely, brain, which has the second-highest per-kilogram metabolic rate in the set, is essentially insulin-independent for glucose uptake. Summing across the six anchors, roughly 41% of accounted resting energy expenditure sits in insulin-responsive (GLUT4) tissue, 26% in insulin-modulated liver, and 34% in insulin-independent tissue. Mitochondrial density, the variable that carried the entire Layer 4 compilation in Section 5, therefore conveys essentially no information about whether a tissue is a target of insulin action.

**Figure 7.**
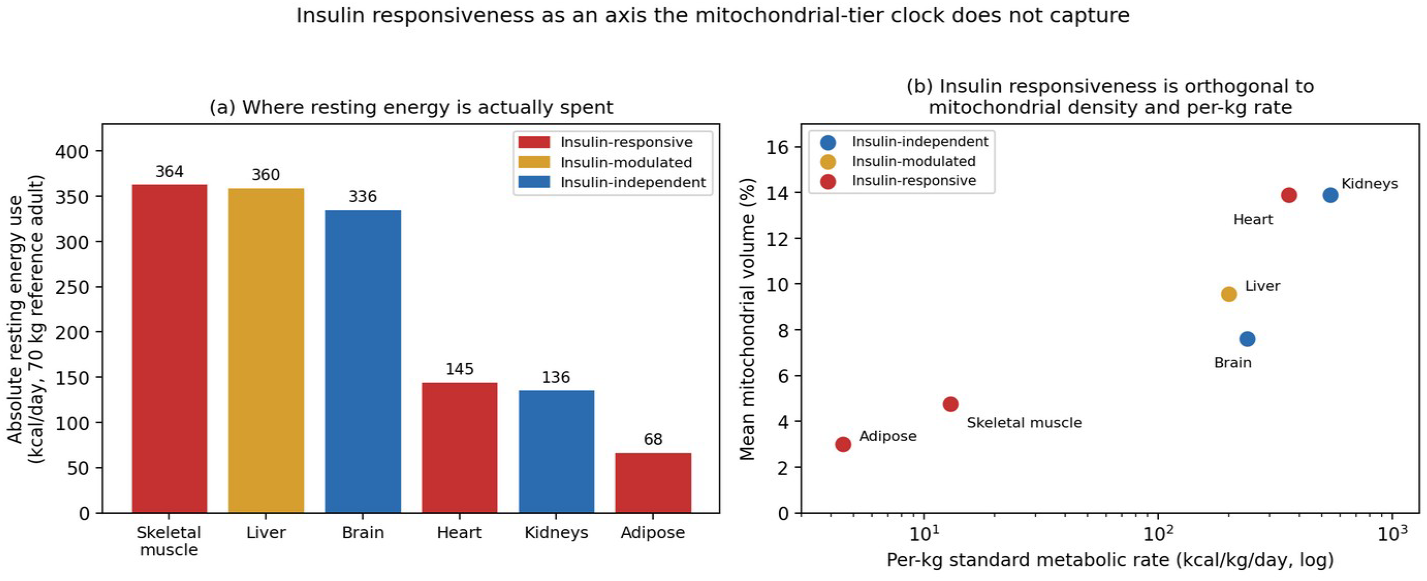
(a) Absolute resting energy expenditure by organ for a 70 kg reference adult (per-kg SMR × organ mass), coloured by insulin-responsiveness class. Skeletal muscle, the lowest-tier tissue per kilogram, is the single largest absolute consumer and the principal insulin target. (b) Mean mitochondrial volume fraction against per-kg SMR for the same organs. Heart and kidney are displaced slightly along the x-axis for visibility; the dataset assigns them identical anchor values (440 kcal/kg/day, 13.88% mitochondrial volume), so they coincide exactly on the underlying data.

### 8.3 Per-kilogram rate versus absolute flux

Table 5 also exposes a scaling issue that Section 5 suppressed by working in per-kilogram units throughout. Skeletal muscle has the second-lowest per-kilogram metabolic rate of any organ in the set (13 kcal·kg^−1^·day^−1^, roughly one-thirtieth that of heart), yet because it constitutes some 40% of body mass it is the single largest absolute consumer of resting energy in the body, at approximately 364 kcal/day — comparable to liver and greater than brain. A framework that ranks organs by mitochondrial density alone, as Figure 3 does, will systematically under-weight exactly the tissue where most insulin-directed glucose actually goes. This is not a flaw in the mitochondrial measurements; it is a consequence of choosing an intensive rather than an extensive variable at the Layer 4 compilation step, and it compounds the cell-number weighting problem already noted as Limit 10.4.

### 8.4 A substrate-selection axis the tier scale collapses

Finally, insulin resistance implicates the dominant-fuel field of the dataset in a way the ordinal tier scale cannot represent. Randle and colleagues’ glucose–fatty-acid cycle described a reciprocal competition in which elevated fatty-acid availability suppresses glucose oxidation and uptake in muscle and adipose tissue [24] — a substrate-selection switch that operates without any change in a cell’s total oxidative capacity. In the terms of Section 4, this is a fuel-pathway relabeling event with no accompanying tier change, and Table 2 records that 78% of lineage edges involve exactly such relabelings. The framework as built treats those relabelings as secondary to tier transitions. Insulin resistance suggests the opposite ordering for adult pathophysiology: the clinically consequential change is which substrate a committed cell preferentially oxidizes, not how much total oxidative machinery it carries. A complete organ clock would need to track fuel selection as a first-class state variable rather than as an annotation attached to tier.

## 9. Machine Learning and Deep Learning Tools Within Reach

Sections 2–8 were deliberately built from auditable, hand-specified formalisms — tree grammars, Shannon entropy, thermodynamic-style aggregation — in the same spirit as the layered genome framework this paper’s structure follows: each step is checkable by inspection, at the cost of leaving most of the available signal in the raw data untouched. A large and rapidly maturing set of machine learning and deep learning tools now exists that could extract more of that signal directly from primary single-cell data, at each layer, without requiring the intermediate hand-built representations of Sections 2–6. This section goes layer by layer through what is concretely within reach today, distinguishing tools that already exist and could be applied immediately from the modeling gaps they would not close.

### 9.1 Layer 1: graph neural networks in place of a hand-assigned grammar

Section 2 classified lineage transitions into grammar classes by inspection (Table 1). A graph neural network — a graph convolutional network [16] or a graph attention network [17] — operating directly on the lineage tree as a graph object could instead learn which local subtree patterns predict a metabolic switch, without an analyst pre-assigning grammar classes. Graph attention in particular is a natural fit for the high-branching hub nodes of Section 2.1: its attention weights would show empirically which of a hub’s many children carry most of the predictive signal for the parent’s eventual metabolic fate, in place of the qualitative claim that such nodes require context-free or context-sensitive rewriting.

### 9.2 Layer 2: learned feature importance in place of binned entropy

The depth-versus-fate variance decomposition in Section 3.2 used a simple, fully auditable calculation (pooled within-group sums of squares) on a five-level ordinal tier. A gradient-boosted tree ensemble trained to predict continuous mitochondrial volume fraction from lineage depth, compartment, germ layer, and branching context, combined with SHAP feature attributions [18], would decompose the same question — how much predictive weight falls on fate identity versus developmental time — without discretizing tier into five bins first, and would additionally reveal any depth-by-compartment interaction effects the current linear variance decomposition cannot detect.

### 9.3 Layer 4: single-cell foundation models in place of an unweighted average

Section 5.1’s organ-level aggregation averaged mitochondrial volume percentage across constituent cell types, unweighted by cell abundance (Limit 10.4). Pretrained single-cell foundation models — Geneformer, pretrained on tens of millions of single-cell transcriptomes to make context-aware predictions in network biology [19]; scGPT, a generative transformer pretrained on over 33 million cells [20]; and scVI, a variational-autoencoder framework for single-cell expression data [21] — could replace that unweighted average with a learned, high-dimensional embedding of each organ’s actual cell-type composition, fine-tuned specifically to predict organ SMR as the downstream task. This would upgrade Figure 3’s five-point consistency check into a genuine held-out prediction problem, addressed directly in Prediction 2 of Section 11.

### 9.4 Layer 5: RNA velocity as an existing, direct test of the clock itself

Of every tool surveyed here, one maps almost exactly onto the question Section 6 asks. RNA velocity estimates the time derivative of a cell’s transcriptional state from the ratio of unspliced to spliced mRNA, recovering a high-dimensional vector that predicts each cell’s near-future state [22]. scVelo, the generalized dynamical-modeling implementation of RNA velocity, explicitly computes a per-cell “latent time” that its own documentation describes as the cell’s internal clock, approximating real developmental time from transcriptional dynamics alone, and separately flags regimes of fate commitment along the trajectory [23]. This is, concretely, a tool that already exists and could be run today to test the commitment-horizon model directly: if scVelo latent time (a continuous, transcriptome-derived proxy for developmental time) predicts metabolic tier no better than lineage depth does in the present analysis, while fate/compartment identity continues to dominate, that would be direct single-cell evidence for the commitment-horizon model rather than the inference from curated, tree-level annotations this paper currently relies on.

### 9.5 What these tools would not resolve

None of this is a substitute for the validation strategies below, and each tool carries a specific limitation worth stating rather than glossing over. Geneformer, scGPT, and scVI are pretrained predominantly on adult and disease-state single-cell atlases, not on lineage-resolved material spanning zygote to terminal fate, so their embeddings may transfer poorly to early developmental states without additional fine-tuning on developmental data this paper does not have. Graph neural network attention weights and foundation-model embeddings are, in exchange for predictive power, considerably less auditable than the hand-specified grammar classes and thermodynamic aggregation of Sections 2 and 5 — the same interpretability-versus-performance tension already flagged for RNAformer versus SCFGs in the sequence-based framework this paper’s structure follows. And RNA velocity’s own literature documents systematic failure modes when its constant-splicing-rate assumption is violated [23], so a null result from scVelo latent time would need to be checked against those known failure modes before being read as evidence against the commitment-horizon model.

### 9.6 Validation strategies not yet run

Unlike sequence-to-circuit compilers such as Cello, which can be run today to test whether a thermodynamic layer composes with an actual DNA construct, none of the ML/DL applications above have been executed against primary data for this paper; Sections 9.1–9.4 describe what is within reach, not what has been done. Three further strategies, independent of any specific ML tool, remain useful complements.

#### Single-cell metabolic flux profiling across pseudotime

Pseudotime-ordered single-cell metabolomic or Seahorse-style flux data, mapped onto a lineage-annotated atlas, would let the depth-versus-fate variance decomposition in Section 3.2 be repeated on primary measurements rather than on curated annotations, directly testing whether the ∼75%/1% split holds outside this dataset.

#### Agent-based lineage simulation

An agent-based model in which simulated progenitors carry a tier state that either drifts continuously with each division (continuous-clock null model) or resets only at designated commitment events (commitment-horizon model) would let the two hypotheses in Section 6 be compared on simulated ground truth before being fit to noisy biological data.

#### Organelle-clearance perturbation

Because the down-shift edges identified in Section 4.2 correspond to known organelle-clearance programs (lens fibre cells, keratinocytes), genetically blocking clearance (e.g., DLAD/PLAAT pathway disruption in lens [3]) and re-measuring mitochondrial volume fraction would test whether the predicted tier decrease is causally tied to the clearance program rather than to an unrelated correlate of terminal differentiation.

## 10. Limits of the Framework

A frank accounting of limits is as important here as the results themselves.

### 10.1 Roughly half the dataset is not directly measured

Of 396 rows, 173 (44%) are labeled “Measured,” while 172 (43%) are “Estimated,” 26 (7%) “Tissue-inferred,” and 25 (6%) “Morphology-inferred” — roughly half the dataset by row count. Every result in Sections 3–6 inherits this uncertainty, and none of the entropy or variance-decomposition numbers should be read as more precise than the underlying annotation basis allows.

### 10.2 The organ-SMR anchor set is small and non-independent

As noted in Section 5.2, the r = 0.90 correlation rests on five organ categories drawn from one classical reference table [4,5], not five independently sourced measurements. It is a consistency check, not a validated predictive model.

### 10.3 The tree is a curated consensus, not a lineage-traced measurement

Human lineage relationships below the level of well-studied model organisms are reconstructed from developmental biology literature rather than directly lineage-traced (as is now possible in mouse or zebrafish with CRISPR-based barcoding). Branch points, especially in less-studied clades, may not reflect the true bifurcation structure.

### 10.4 No cell-number weighting is available

The organ-level aggregation in Section 5 averages mitochondrial volume fraction across constituent cell types without weighting by their actual relative abundance in the tissue, which is not present in this dataset. A neuron-dominated versus glia-dominated weighting of brain tissue, for instance, could shift the brain data point in Figure 3 substantially.

### 10.5 A five-level ordinal tier is a coarse discretization

The entropy analysis in Section 3.1 is sensitive to the number of tiers chosen; collapsing or expanding the five-level scale would change the absolute entropy values, though the qualitative depth-versus-fate contrast in Section 3.2, which does not depend on the entropy calculation, would likely be more robust to this choice.

### 10.6 Composability across layers is not demonstrated end-to-end

As in any layered decomposition, showing that each layer individually contains real signal (Sections 2–6) is not the same as showing that the layers compose into a single predictive pipeline from raw lineage topology to organ SMR. Section 9 describes what that end-to-end test would require; it has not been run here.

### 10.7 The cross-species comparison in Section 7 is illustrative, not a fresh comparative analysis

The ten species in Table 4 are a hand-selected, literature-sourced set chosen to include well-documented longevity outliers, not a random or phylogenetically representative sample. Body masses and observed maximum lifespans are drawn from standard secondary sources rather than re-derived from primary data, and breathing rates are independently measured for only four of the ten species, with the remaining six shown as equation-evaluated points that carry no evidential weight (Section 7.1, Figure 5); an earlier version of this figure plotted the equation’s output for all ten species and was therefore circular. The analysis is also endotherm-only, which excludes the ectothermic longevity records where the rate-of-living hypothesis appears to hold (Section 7.5). The qualitative point (four species living several-fold longer than mass alone predicts, for documented non-pace-of-living reasons) is well supported by the primary literature cited; the specific numbers in Table 4 should be treated as representative rather than exact.

### 10.8 The machine learning tools of Section 9 have not been run on this problem

Geneformer, scGPT, scVI, and scVelo are real, published, and usable today, but none has been applied to this dataset or to a lineage-resolved developmental atlas with matched metabolic phenotypes. Their pretraining corpora skew toward adult and disease-state tissue, so their out-of-the-box performance on zygote-to-terminal developmental data is unknown and could be poor without fine-tuning on data this paper does not have access to. Section 9 should be read as a map of what is reachable, not a report of results obtained.

### 10.9 The insulin overlay of Section 8 is literature-derived, not dataset-derived

The cell-lineage dataset contains no insulin-signalling, GLUT4-expression, or glucose-uptake fields. Every transporter and insulin-class assignment in Table 5 was mapped onto the organ anchors by hand from the insulin-physiology literature [25–28], and the organ masses used to convert per-kilogram rates into absolute energy flux are standard reference-adult values rather than measurements matched to the same source as the SMR anchors. The dissociation reported in Section 8.2 is therefore a juxtaposition of two independently sourced literatures, which is sufficient to show that mitochondrial density does not predict insulin responsiveness, but is not a within-dataset statistical result and carries no p-value.

## 11. Conclusion: Testable Predictions and Falsification Conditions

The value of the framework proposed here rests on whether it generates predictions a purely descriptive cell atlas does not. We close with eight, each paired with a stated falsification condition.

### (1) Fate identity dominates over developmental time in independent data

Compartment/fate identity should continue to explain the large majority of metabolic-tier variance when the analysis of Section 3.2 is repeated on an independently collected single-cell metabolic atlas.

*Falsification: an independent atlas shows lineage depth or pseudotime explaining a comparable or larger share of variance than fate identity*.

### (2) The mitochondrial-fraction / SMR relationship generalizes out of sample

The occupancy-style aggregation in Section 5.1 should predict the SMR ordering of organs not among the five original Elia/Wang anchor categories (e.g., gastrointestinal tract, lung, spleen) once those organs’ constituent cell types are annotated.

*Falsification: newly annotated organs show mitochondrial-fraction rankings inconsistent with their measured SMR ordering*.

### (3) Down-shift edges are enriched for organelle-clearance programs

Tier-decreasing lineage edges, considered as a class, should be enriched for terminal-differentiation programs involving active organelle clearance (lens fibre cells, keratinocytes, reticulocytes) relative to tier-stable or tier-increasing edges.

*Falsification: down-shift edges show no enrichment for organelle-clearance biology relative to other edge types*.

### (4) The commitment-horizon model out-predicts a continuous depth model. (Partially tested; see Section 6.4.)

Run on the present dataset, the nearest-ancestor predictor beat the null model (p = 0.015) and had lower point-estimate error than the depth regression, but the difference between them was not significant (ΔMAE = −0.042, 95% CI −0.098 to +0.012, p = 0.175). The prediction therefore stands as untested in its discriminating form and requires primary single-cell data with a finer metabolic readout than a five-level ordinal tier.

*Falsification: with adequately powered data and a continuous metabolic readout, a depth-indexed regression performs equally well or better than the commitment-horizon predictor on held-out cells*.

### (5) Branching multiplicity and metabolic-switch frequency remain uncorrelated

The near-zero correlation reported in Section 4.2 (r = −0.015) between a node’s branching factor and its outgoing switch fraction should replicate in independently curated lineage datasets, supporting the view that fate specification and metabolic switching are separable developmental programs.

*Falsification: independent datasets show a substantial positive or negative correlation between branching factor and switch frequency*.

### (6) Longevity-outlier species show stronger within-species canalization of metabolic fate, not a lower pace of living

Extending the depth-versus-fate variance decomposition of Section 3.2 to single-cell lineage data from a high-LQ species (e.g., naked mole rat) should show fate identity explaining an even larger share of cellular metabolic-tier variance than the ∼75% observed in humans, consistent with these species achieving longevity through tighter metabolic canalization/damage suppression rather than through slower cellular pace — the same distinction Section 7.3 draws at the organism level.

*Falsification: a longevity-outlier species shows a depth-versus-fate split no different from, or weaker than, the human result, indicating its longevity advantage operates through a mechanism unrelated to within-lineage metabolic canalization*.

### (7) scVelo latent time reproduces the depth-versus-fate dissociation on real single-cell data

Applying scVelo [23] to a lineage-resolved single-cell RNA-seq atlas and regressing metabolic-tier proxies (e.g., mitochondrial gene expression score) on latent time versus on cell-type/compartment identity should reproduce the qualitative result of Section 3.2: fate identity explaining substantially more variance than the continuous, transcriptome-derived time axis.

*Falsification: scVelo latent time explains a comparable or larger share of metabolic-marker variance than fate identity does, favoring a continuous-clock model over the commitment-horizon model on real transcriptomic dynamics rather than on the curated tree used here*.

### (8) Insulin responsiveness is uncorrelated with mitochondrial density across tissues

Extending the overlay of Section 8.2 to a larger panel of human tissues with matched measurements — insulin-stimulated glucose uptake by PET or clamp-with-catheterization on one axis, mitochondrial volume fraction or OXPHOS protein abundance on the other — should show no systematic relationship between the two, confirming that hormonal responsiveness and resting oxidative capacity are separable tissue properties.

*Falsification: a matched multi-tissue panel shows insulin-stimulated glucose disposal scaling with mitochondrial content, indicating that the Layer 4 mitochondrial variable already captures insulin responsiveness and that no separate substrate-selection axis is required*.

None of these predictions requires the metabolic tier of a cell to be literally computed the way a program is executed. They require only that treating lineage as a computable structure — a tree grammar compiled, via a small number of discrete switch events, into an emergent organ-level energetic network — generates checkable claims that a purely correlational cell atlas, read as a static catalogue, does not.

## Methods and Data Note

Sections 2–6 and their figures/tables were computed directly from a single curated dataset of 396 human cell types spanning zygote to terminal identity, with fields for parent identity, germ layer, anatomical compartment, lineage depth, terminal status, five-level metabolic tier (T0–T5), mitochondrial volume percentage (parsed from point estimates and ranges by taking the arithmetic midpoint), dominant fuel/pathway, organ-level SMR where available, an evidentiary basis flag (Measured / Estimated / Tissue-inferred / Morphology-inferred), free-text notes, and full lineage path. Correlations are Pearson product-moment correlations on the numeric tier scale (T0=0 … T5=5) unless otherwise noted; entropy is Shannon entropy in bits; the between/within-compartment variance decomposition pools within-group sums of squares across all 227 represented compartments. Section 7 and its figures/table draw instead on external comparative-biology literature: body masses and observed maximum lifespans are representative values compiled from standard secondary sources; breathing rates are computed from Stahl’s allometric equation [9]; predicted maximum lifespans and longevity quotients are computed from the allometric equation of de Magalhães et al. [10]. All code and the full data table are available on request.

### Dataset Provenance

The analyses in Sections 2–6 rest entirely on one curated table of 396 human cell types, and its provenance therefore determines how much weight the results can carry. The table was supplied to the authors as a single compiled file rather than generated by the authors from primary literature, and it records, for each cell type, a parent identifier, germ layer, anatomical compartment, lineage depth, terminal status, a five-level ordinal metabolic tier, mitochondrial volume percentage, dominant fuel/pathway, organ-level SMR where applicable, a free-text note, the full lineage path, and an evidentiary basis flag.

That basis flag is the only per-row provenance information the file carries, and it is coarse: 173 rows (44%) are marked Measured, 172 (43%) Estimated, 26 (7%) Tissue-inferred and 25 (6%) Morphology-inferred. Critically, the file does not record a primary citation for any individual row. We can therefore state which rows the compiler regarded as measured, but not what they were measured by, in what preparation, or by whom. Readers should treat the dataset as an expert-curated consensus compilation of the developmental and cell-biological literature, of the kind routinely used to scaffold hypotheses, and not as a primary measurement resource. Two checks partially offset this. The organ-level SMR values reproduce the classical Elia/Wang reference figures exactly [4,5], confirming that at least that field was drawn from the standard literature. And the mitochondrial-volume field independently recovers the known between-organ ordering of metabolic rate (Section 5.1), which a randomly or carelessly assembled table would not. Neither check substitutes for row-level citation, and establishing that would be the single highest-value improvement to this work.

## Data Availability

No new experimental data were generated for this work. The cell-lineage-metabolism table analysed here, together with a single self-contained analysis script that regenerates every number, table and figure in this paper from that table — the variance decomposition of Section 3.2, both regressions of Tables 1b and 2b, the lineage-edge switch counts of Section 4, the entropy curve of Figure 2, the five-organ fit of Section 5.1, and the full cross-validation of Section 6.4 including bootstrap confidence intervals and paired Wilcoxon tests — are deposited at [REPOSITORY DOI TO BE INSERTED ON SUBMISSION]. The script has no dependencies beyond pandas, numpy, scipy, scikit-learn and matplotlib, and reproduces every reported value deterministically under fixed random seeds. The insulin-responsiveness assignments of Section 8 and the comparative-longevity values of Section 7 are literature-derived and are cited individually rather than deposited.

## Author Contributions

A.M.S. and L.P. conceived the framework. A.M.S. performed the computational analyses and prepared the figures. A.M.S. and L.P. wrote the manuscript. Both authors read and approved the final version.

## Acknowledgments

The authors thank colleagues at Ainnocence Inc. for discussions and feedback. We acknowledge the developmental biology and comparative physiology communities whose published cell-atlas and longevity data underpin every estimate in this paper.

